## Supplemental Files for "*couple*CoC+: an information-theoretic co-clustering-based transfer learning framework for the integrative analysis of single-cell genomic data"

### Supplementary Materials

#### 1. Supplementary Text

##### 1.1 *couple*CoC+ algorithm

We first reformulate the loss in mutual information into the form of KL divergence [1,2]. More specifically, for the source data S,

$$\ell_{\mathrm{S}}(C_X, C_Z) = D_{\mathrm{KL}}(p_{\mathrm{S}}(X, Z_{\mathrm{S}})||p^*_{\mathrm{S}}(X, Z_{\mathrm{S}})), \tag{S.1}$$

where $p^*_{\mathrm{S}}(X, Z_{\mathrm{S}})$ is defined as

$$p^*_{\mathrm{S}}(X = x, Z_{\mathrm{S}} = z) = p_{\mathrm{S}}(\tilde{X} = C_X(x), \tilde{Z}_{\mathrm{S}} = C_Z(z)) * \frac{p_{\mathrm{S}}(X = x)}{p_{\mathrm{S}}(\tilde{X} = C_X(x))} \frac{p_{\mathrm{S}}(Z_{\mathrm{S}} = z)}{p_{\mathrm{S}}(\tilde{Z}_{\mathrm{S}} = C_Z(z))}. \tag{S.2}$$

Further, we have

$$\begin{aligned} D_{\mathrm{KL}}(p_{\mathrm{S}}(X, Z_{\mathrm{S}})||p^*_{\mathrm{S}}(X, Z_{\mathrm{S}})) &= \sum_{i=1}^{N_{\mathrm{S}}} \sum_{x \in \{x: C_X(x)=i\}} p_{\mathrm{S}}(X = x) D_{\mathrm{KL}}(p_{\mathrm{S}}(Z_{\mathrm{S}}|X = x)||p^*_{\mathrm{S}}(Z_{\mathrm{S}}|\tilde{X} = i, X = x)) \\ &= \sum_{j=1}^{K} \sum_{z \in \{z: C_Z(z)=j\}} p_{\mathrm{S}}(Z_{\mathrm{S}} = z) D_{\mathrm{KL}}(p_{\mathrm{S}}(X|Z_{\mathrm{S}} = z)||p^*_{\mathrm{S}}(X|\tilde{Z}_{\mathrm{S}} = j, Z_{\mathrm{S}} = z)), \end{aligned} \tag{S.3}$$

where $p^*_{\mathrm{S}}(Z_{\mathrm{S}} = z|\tilde{X} = i, X = x) \triangleq \frac{p^*_{\mathrm{S}}(X=x, Z_{\mathrm{S}}=z)}{p_{\mathrm{S}}(X=x)}$, $z = 1, \ldots, q$, for any $x \in \{x : C_X(x) = i\}$, and $p^*_{\mathrm{S}}(X = x|\tilde{Z}_{\mathrm{S}} = j, Z_{\mathrm{S}} = z) \triangleq \frac{p^*_{\mathrm{S}}(X=x, Z_{\mathrm{S}}=z)}{p_{\mathrm{S}}(Z_{\mathrm{S}}=z)}$, $x = 1, \ldots, n_{\mathrm{S}}$, for any $z \in \{z : C_Z(z) = j\}$. Details on the derivation of formulas (S.1) and (S.3) are presented in [1] and [2]. For the target data T and U, we can rewrite $\ell_{\mathrm{T}}$ and $\ell_{\mathrm{U}}$ similarly as we rewrote $\ell_{\mathrm{S}}$ in formulas (S.1) and (S.3) as follows:

$$\ell_{\mathrm{T}}(C_Y, C_Z) = D_{\mathrm{KL}}(p_{\mathrm{T}}(Y_{\mathrm{T}}, Z_{\mathrm{T}})||p^*_{\mathrm{T}}(Y_{\mathrm{T}}, Z_{\mathrm{T}})), \tag{S.4}$$

and

$$\begin{aligned} D_{\mathrm{KL}}(p_{\mathrm{T}}(Y_{\mathrm{T}}, Z_{\mathrm{T}})||p^*_{\mathrm{T}}(Y_{\mathrm{T}}, Z_{\mathrm{T}})) &= \sum_{i=1}^{N_{\mathrm{T}}} \sum_{y \in \{y: C_Y(y)=i\}} p_{\mathrm{T}}(Y_{\mathrm{T}} = y) D_{\mathrm{KL}}(p_{\mathrm{T}}(Z_{\mathrm{T}}|Y_{\mathrm{T}} = y)||p^*_{\mathrm{T}}(Z_{\mathrm{T}}|\tilde{Y}_{\mathrm{T}} = i, Y_{\mathrm{T}} = y)) \\ &= \sum_{j=1}^{K} \sum_{z \in \{z: C_Z(z)=j\}} p_{\mathrm{T}}(Z_{\mathrm{T}} = z) D_{\mathrm{KL}}(p_{\mathrm{T}}(Y_{\mathrm{T}}|Z_{\mathrm{T}} = z)||p^*_{\mathrm{T}}(Y_{\mathrm{T}}|\tilde{Z}_{\mathrm{T}} = j, Z_{\mathrm{T}} = z)), \end{aligned} \tag{S.5}$$

and

$$\ell_{\mathrm{U}}(C_Y, C_U) = D_{\mathrm{KL}}(p_{\mathrm{U}}(Y_{\mathrm{U}}, Z_{\mathrm{U}})||p^*_{\mathrm{U}}(Y_{\mathrm{U}}, Z_{\mathrm{U}})), \tag{S.6}$$

and

$$\begin{aligned} D_{\mathrm{KL}}(p_{\mathrm{U}}(Y_{\mathrm{U}}, Z_{\mathrm{U}})||p^*_{\mathrm{U}}(Y_{\mathrm{U}}, Z_{\mathrm{U}})) &= \sum_{i=1}^{N_{\mathrm{T}}} \sum_{y \in \{y: C_Y(y)=i\}} p_{\mathrm{U}}(Y_{\mathrm{U}} = y) D_{\mathrm{KL}}(p_{\mathrm{U}}(Z_{\mathrm{U}}|Y_{\mathrm{U}} = y)||p^*_{\mathrm{U}}(Z_{\mathrm{U}}|\tilde{Y}_{\mathrm{U}} = i, Y_{\mathrm{U}} = y)) \\ &= \sum_{j=1}^{K_0} \sum_{u \in \{u: C_U(u)=j\}} p_{\mathrm{U}}(Z_{\mathrm{U}} = u) D_{\mathrm{KL}}(p_{\mathrm{U}}(Y_{\mathrm{U}}|Z_{\mathrm{U}} = u)||p^*_{\mathrm{U}}(Y_{\mathrm{U}}|\tilde{Z}_{\mathrm{U}} = j, Z_{\mathrm{U}} = u)). \end{aligned} \tag{S.7}$$

(Note that $K_0$ in formula (S.7) denotes the number of feature clusters in data U.) Therefore, the following optimization problem

$$\begin{aligned} \underset{\substack{C_Y, C_X, C_Z, C_U \\ h_{\mathrm{T},N_{\mathrm{sub}}}, h_{\mathrm{S},N_{\mathrm{sub}}}}}{\operatorname{argmin}} \quad & \ell_{\mathrm{T}}(C_Y, C_Z) + \lambda \ell_{\mathrm{S}}(C_X, C_Z) + \beta \ell_{\mathrm{U}}(C_Y, C_U) \\ & + \gamma D_{\mathrm{KL}}(\hat{p}_{\mathrm{T}}(\tilde{Y}_{h_{\mathrm{T},N_{\mathrm{sub}}}}, \tilde{Z}_{\mathrm{T}})||\hat{p}_{\mathrm{S}}(\tilde{X}_{h_{\mathrm{S},N_{\mathrm{sub}}}}, \tilde{Z}_{\mathrm{S}})). \end{aligned} \tag{S.8}$$

can be rewritten as:

$$\begin{aligned} \underset{\substack{C_Y, C_X, C_Z, C_U \\ h_{\mathrm{T},N_{\mathrm{sub}}}, h_{\mathrm{S},N_{\mathrm{sub}}}}}{\operatorname{argmin}} \quad & D_{\mathrm{KL}}(p_{\mathrm{S}}(X, Z_{\mathrm{S}})||p^*_{\mathrm{S}}(X, Z_{\mathrm{S}})) + \lambda D_{\mathrm{KL}}(p_{\mathrm{T}}(Y_{\mathrm{T}}, Z_{\mathrm{T}})||p^*_{\mathrm{T}}(Y_{\mathrm{T}}, Z_{\mathrm{T}})) + \beta D_{\mathrm{KL}}(p_{\mathrm{U}}(Y_{\mathrm{U}}, Z_{\mathrm{U}})||p^*_{\mathrm{U}}(Y_{\mathrm{U}}, Z_{\mathrm{U}})) \\ & + \gamma D_{\mathrm{KL}}(\hat{p}_{\mathrm{T}}(\tilde{Y}_{h_{\mathrm{T},N_{\mathrm{sub}}}}, \tilde{Z}_{\mathrm{T}})||\hat{p}_{\mathrm{S}}(\tilde{X}_{h_{\mathrm{S},N_{\mathrm{sub}}}}, \tilde{Z}_{\mathrm{S}})) \end{aligned} \tag{S.9}$$

Next, the optimization problem (S.9) can be solved by iteratively updating $C_Y$, $C_X$, $C_Z$, $C_U$, $h_{\mathrm{T},N_{\mathrm{sub}}}$ and $h_{\mathrm{S},N_{\mathrm{sub}}}$ as follows:

- Given $C_X$, $C_Z$, $C_U$, $h_{\mathrm{T},N_{\mathrm{sub}}}$ and $h_{\mathrm{S},N_{\mathrm{sub}}}$, update $C_Y$. The optimization problem (S.9) is equivalent to minimizing

  $$\sum_{i=1}^{N_{\mathrm{T}}} \sum_{y \in \{y: C_Y(y)=i\}} p_{\mathrm{T}}(Y_{\mathrm{T}} = y) Q(\tilde{Y}_{\mathrm{T}} = i, Y_{\mathrm{T}} = y | C_X, C_Z, C_U, h_{\mathrm{S},N_{\mathrm{sub}}}, h_{\mathrm{T},N_{\mathrm{sub}}}),$$

  where

  $$\begin{aligned} & Q(\tilde{Y}_{\mathrm{T}} = i, Y_{\mathrm{T}} = y | C_X, C_Z, C_U, h_{\mathrm{S},N_{\mathrm{sub}}}, h_{\mathrm{T},N_{\mathrm{sub}}}) \triangleq D_{\mathrm{KL}}(p_{\mathrm{T}}(Z_{\mathrm{T}}|Y_{\mathrm{T}} = y)||p^*_{\mathrm{T}}(Z_{\mathrm{T}}|\tilde{Y}_{\mathrm{T}} = i, Y_{\mathrm{T}} = y)) + \\ & \beta \frac{p_{\mathrm{U}}(Y_{\mathrm{U}} = y)}{p_{\mathrm{T}}(Y_{\mathrm{T}} = y)} D_{\mathrm{KL}}(p_{\mathrm{U}}(Z_{\mathrm{U}}|Y_{\mathrm{U}} = y)||p^*_{\mathrm{U}}(Z_{\mathrm{U}}|\tilde{Y}_{\mathrm{U}} = i, Y_{\mathrm{U}} = y)) + \frac{\gamma D_{\mathrm{KL}}(\hat{p}_{\mathrm{T}}(\tilde{Y}_{h_{\mathrm{T},N_{\mathrm{sub}}}}, \tilde{Z}_{\mathrm{T}})||\hat{p}_{\mathrm{S}}(\tilde{X}_{h_{\mathrm{S},N_{\mathrm{sub}}}}, \tilde{Z}_{\mathrm{S}}))}{n_{\mathrm{T}} p_{\mathrm{T}}(Y = y)}. \end{aligned}$$

  We iteratively update the cluster assignment $C_Y(y)$ for each cell $y$ ($y = 1, \ldots, n_{\mathrm{T}}$) in the target data, fixing the cluster assignment for the other cells:

  $$C_Y(y) = \underset{i \in \{1, \ldots, N_{\mathrm{T}}\}}{\operatorname{argmin}} Q(\tilde{Y}_{\mathrm{T}} = i, Y_{\mathrm{T}} = y | C_X, C_Z, C_U, h_{\mathrm{S},N_{\mathrm{sub}}}, h_{\mathrm{T},N_{\mathrm{sub}}}). \tag{S.10}$$

- Given $C_Y$, $C_Z$, $C_U$, $h_{\mathrm{T},N_{\mathrm{sub}}}$ and $h_{\mathrm{S},N_{\mathrm{sub}}}$, update $C_X$. The optimization problem (S.9) is equivalent to minimizing

  $$\sum_{j=1}^{N_{\mathrm{S}}} \sum_{x \in \{x: C_X(x)=j\}} p_{\mathrm{S}}(X = x) Q(\tilde{X} = j, X = x | C_Y, C_Z, C_U, h_{\mathrm{S},N_{\mathrm{sub}}}, h_{\mathrm{T},N_{\mathrm{sub}}}),$$

  where

  $$\begin{aligned} & Q(\tilde{X} = j, X = x | C_Y, C_Z, C_U, h_{\mathrm{S},N_{\mathrm{sub}}}, h_{\mathrm{T},N_{\mathrm{sub}}}) \\ & \triangleq \lambda D_{\mathrm{KL}}(p_{\mathrm{S}}(Z_{\mathrm{S}}|X = x)||p^*_{\mathrm{S}}(Z_{\mathrm{S}}|\tilde{X} = j, X = x)) + \frac{\gamma D_{\mathrm{KL}}(\hat{p}_{\mathrm{T}}(\tilde{Y}_{h_{\mathrm{T},N_{\mathrm{sub}}}}, \tilde{Z}_{\mathrm{T}})||\hat{p}_{\mathrm{S}}(\tilde{X}_{h_{\mathrm{S},N_{\mathrm{sub}}}}, \tilde{Z}_{\mathrm{S}}))}{n_{\mathrm{S}} p_{\mathrm{S}}(X = x)}. \end{aligned}$$

We iteratively update the cluster assignment $C_X(x)$ for each cell $x$ $(x = 1, \ldots, n_{\mathrm{S}})$ in the source data, fixing the cluster assignment for the other cells:

$$C_X(x) = \underset{j\in\{1,\ldots,N_{\mathrm{S}}\}}{\operatorname{argmin}} Q(\tilde{X} = j, X = x | C_Y, C_Z, C_U, h_{\mathrm{S},N_{\mathrm{sub}}}, h_{\mathrm{T},N_{\mathrm{sub}}}). \tag{S.11}$$

- Given $C_X$, $C_Y$, $C_U$, $h_{\mathrm{T},N_{\mathrm{sub}}}$ and $h_{\mathrm{S},N_{\mathrm{sub}}}$, update $C_Z$. The optimization problem (S.9) is equivalent to minimizing

$$\sum_{s=1}^{K} \sum_{z\in\{z:\tilde{Z}(z)=s\}} R(\tilde{Z}_{\mathrm{T}} = s, Z_{\mathrm{T}} = z | C_Y, C_X, C_U, h_{\mathrm{S},N_{\mathrm{sub}}}, h_{\mathrm{T},N_{\mathrm{sub}}}),$$

where

$$R(\tilde{Z}_{\mathrm{T}} = s, Z_{\mathrm{T}} = z | C_Y, C_X, C_U, h_{\mathrm{S},N_{\mathrm{sub}}}, h_{\mathrm{T},N_{\mathrm{sub}}}) \triangleq p_{\mathrm{T}}(Z_{\mathrm{T}} = z) D_{\mathrm{KL}}(p_{\mathrm{T}}(Y_{\mathrm{T}}|Z_{\mathrm{T}} = z)||p^*_{\mathrm{T}}(Y_{\mathrm{T}}|\tilde{Z}_{\mathrm{T}} = s, Z_{\mathrm{T}} = z))$$
$$+ \lambda p_{\mathrm{S}}(Z_{\mathrm{S}} = z) D_{\mathrm{KL}}(p_{\mathrm{S}}(X|Z_{\mathrm{S}} = z)||p^*_{\mathrm{S}}(X|\tilde{Z}_{\mathrm{S}} = s, Z_{\mathrm{S}} = z)) + \frac{\gamma D_{\mathrm{KL}}(\hat{p}_{\mathrm{T}}(\tilde{Y}_{h_{\mathrm{T},N_{\mathrm{sub}}}}, \tilde{Z}_{\mathrm{T}})||\hat{p}_{\mathrm{S}}(\tilde{X}_{h_{\mathrm{S},N_{\mathrm{sub}}}}, \tilde{Z}_{\mathrm{S}}))}{q}.$$

We iteratively update the cluster assignment $C_Z(z)$ for each feature $z$ $(z = 1, \ldots, q)$, fixing the cluster assignment for the other features:

$$C_Z(z) = \underset{s\in\{1,\ldots,K\}}{\operatorname{argmin}} R(\tilde{Z}_{\mathrm{T}} = s, Z_{\mathrm{T}} = z | C_Y, C_X, C_U, h_{\mathrm{S},N_{\mathrm{sub}}}, h_{\mathrm{T},N_{\mathrm{sub}}}). \tag{S.12}$$

- Given $C_Y$, update $C_U$. The optimization problem (S.9) is equivalent to minimizing

$$\sum_{s=1}^{K_0} \sum_{u\in\{u:\tilde{Z}(u)=s\}} p_{\mathrm{U}}(Z_{\mathrm{U}} = u) D_{\mathrm{KL}}(p_{\mathrm{U}}(Y_{\mathrm{U}}|Z_{\mathrm{U}} = u)||p^*_{\mathrm{U}}(Y_{\mathrm{U}}|\tilde{Z}_{\mathrm{U}} = s, Z_{\mathrm{U}} = u)).$$

We iteratively update the cluster assignment $C_U(u)$ for each feature $u$ $(u = 1, \ldots, q_0)$(Note that $q_0$ denotes the number of features in data U.), fixing the cluster assignment for the other features:

$$C_U(u) = \underset{s\in\{1,\ldots,K_0\}}{\operatorname{argmin}} p_{\mathrm{U}}(Z_{\mathrm{U}} = u) D_{\mathrm{KL}}(p_{\mathrm{U}}(Y_{\mathrm{U}}|Z_{\mathrm{U}} = u)||p^*_{\mathrm{U}}(Y_{\mathrm{U}}|\tilde{Z}_{\mathrm{U}} = s)). \tag{S.13}$$

- Given $C_X$, $C_Y$, $C_Z$, $C_U$, update $h_{\mathrm{T},N_{\mathrm{sub}}}$ and $h_{\mathrm{S},N_{\mathrm{sub}}}$. The optimization problem (S.9) is equivalent to minimizing $D_{\mathrm{KL}}(\hat{p}_{\mathrm{T}}(\tilde{Y}_{h_{\mathrm{T},N_{\mathrm{sub}}}}, \tilde{Z}_{\mathrm{T}})||\hat{p}_{\mathrm{S}}(\tilde{X}_{h_{\mathrm{S},N_{\mathrm{sub}}}}, \tilde{Z}_{\mathrm{S}}))$. So, we obtain the optimal combinations by

$$(h_{\mathrm{T},N_{\mathrm{sub}}}, h_{\mathrm{S},N_{\mathrm{sub}}}) = \underset{\substack{h_{\mathrm{T},N_{\mathrm{sub}}} \\ h_{\mathrm{S},N_{\mathrm{sub}}}}}{\operatorname{argmin}} D_{\mathrm{KL}}(\hat{p}_{\mathrm{T}}(\tilde{Y}_{h_{\mathrm{T},N_{\mathrm{sub}}}}, \tilde{Z}_{\mathrm{T}})||\hat{p}_{\mathrm{S}}(\tilde{X}_{h_{\mathrm{S},N_{\mathrm{sub}}}}, \tilde{Z}_{\mathrm{S}})). \tag{S.14}$$

As mentioned in the main text, $h_{\mathrm{T},N_{\mathrm{sub}}}$ is a permutation of size $N_{\mathrm{sub}}$ for the indexes of the cell clusters in target data, and there are $\frac{N_{\mathrm{T}}!}{N_{\mathrm{sub}}!(N_{\mathrm{T}}-N_{\mathrm{sub}})!}$ total combinations of $h_{\mathrm{T},N_{\mathrm{sub}}}$. $h_{\mathrm{S},N_{\mathrm{sub}}}$ is an ordered permutation of size $N_{\mathrm{sub}}$ for the indexes of the cell clusters in source data, and there are $\frac{N_{\mathrm{S}}!}{(N_{\mathrm{S}}-N_{\mathrm{sub}})!}$ total combinations of $h_{\mathrm{S},N_{\mathrm{sub}}}$. We first calculate the distributions $\tilde{p}_{\mathrm{T}}(\tilde{Y}_{\mathrm{T}}, \tilde{Z}_{\mathrm{T}})$ and $\tilde{p}_{\mathrm{S}}(\tilde{X}, \tilde{Z}_{\mathrm{S}})$ based on $C_Y$, $C_X$ and Equation (1) in the main text (the expression of $\tilde{p}_{\mathrm{T}}(\tilde{Y}_{\mathrm{T}}, \tilde{Z}_{\mathrm{T}})$ is similar to Equation (1)). The rows in the these two matrices are then normalized so the sum of each row equals to one, which reduces the bias of the differences in the sizes of the clusters in the source data S and the data T. We extract the two submatrices $\tilde{p}_{\mathrm{T}}(\tilde{Y}_{h_{\mathrm{T},N_{\mathrm{sub}}}}, \tilde{Z}_{\mathrm{T}})$ and $\tilde{p}_{\mathrm{S}}(\tilde{X}_{h_{\mathrm{S},N_{\mathrm{sub}}}}, \tilde{Z}_{\mathrm{S}})$. Both can be interpreted as the low dimension representations of the subsets of cell clusters, which are further scaled to have total sums equal to 1, and we then obtain the $\hat{p}_{\mathrm{T}}(\tilde{Y}_{h_{\mathrm{T},N_{\mathrm{sub}}}}, \tilde{Z}_{\mathrm{T}})$ and $\hat{p}_{\mathrm{T}}(\tilde{Y}_{h_{\mathrm{S},N_{\mathrm{sub}}}}, \tilde{Z}_{\mathrm{S}})$. Finally, we calculate the KL divergence between the normalized submatrices $\hat{p}_{\mathrm{T}}(\tilde{Y}_{h_{\mathrm{T},N_{\mathrm{sub}}}}, \tilde{Z}_{\mathrm{T}})$ and $\hat{p}_{\mathrm{S}}(\tilde{X}_{h_{\mathrm{S},N_{\mathrm{sub}}}}, \tilde{Z}_{\mathrm{S}})$.

##### 1.2 Summary of the *couple*CoC+ algorithm

To summarize, the procedures of the *couple*CoC+ algorithm are as follows:

1. Initialization. Calculate $p_{\mathrm{S}}(X, Z_{\mathrm{S}})$, $p_{\mathrm{T}}(Y_{\mathrm{T}}, Z_{\mathrm{T}})$ and $p_{\mathrm{U}}(Y_{\mathrm{U}}, Z_{\mathrm{U}})$ using the datasets S, T and U. Initialize the clustering functions $C_Y^{[0]}$, $C_X^{[0]}$, $C_Z^{[0]}$ and $C_U^{[0]}$. Initialize $p_{\mathrm{T}}^{*[0]}(Y_{\mathrm{T}}, Z_{\mathrm{T}})$ and $p_{\mathrm{S}}^{*[0]}(X, Z_{\mathrm{S}})$. Initialize $h_{\mathrm{T},N_{\mathrm{sub}}}^{[0]}$ and $h_{\mathrm{S},N_{\mathrm{sub}}}^{[0]}$.

2. Iterate steps (a) and (b) until convergence.
(a). Fix $p_{\mathrm{T}}^{*[t-1]}(Y_{\mathrm{T}}, Z_{\mathrm{T}})$, $p_{\mathrm{S}}^{*[t-1]}(X, Z_{\mathrm{S}})$, $p_{\mathrm{U}}^{*[t-1]}(Y_{\mathrm{U}}, Z_{\mathrm{U}})$, $h_{\mathrm{T},N_{\mathrm{sub}}}^{[t-1]}$ and $h_{\mathrm{S},N_{\mathrm{sub}}}^{[t-1]}$, and sequentially update $C_Y^{[t]}$, $C_X^{[t]}$, $C_Z^{[t]}$, $C_U^{[t]}$, $h_{\mathrm{T},N_{\mathrm{sub}}}^{[t]}$ and $h_{\mathrm{S},N_{\mathrm{sub}}}^{[t]}$ based on the Equations (S.10), (S.11), (S.12), (S.13) and (S.14).
(b). Fix $C_Y^{[t]}$, $C_X^{[t]}$, $C_Z^{[t]}$, $C_U^{[t]}$, $h_{\mathrm{T},N_{\mathrm{sub}}}^{[t]}$ and $h_{\mathrm{S},N_{\mathrm{sub}}}^{[t]}$, and update $p_{\mathrm{T}}^{*[t]}(Y_{\mathrm{T}}, Z_{\mathrm{T}})$, $p_{\mathrm{S}}^{*[t]}(X, Z_{\mathrm{S}})$ and $p_{\mathrm{U}}^{*[t]}(Y_{\mathrm{U}}, Z_{\mathrm{U}})$.

3. Output the clustering results $C_Y$, $C_X$, $C_Z$, $C_U$, $h_{\mathrm{T},N_{\mathrm{sub}}}$ and $h_{\mathrm{S},N_{\mathrm{sub}}}$ in the last iteration.

##### 1.3 Selecting $N_{\mathrm{sub}}$

We use the following criteria to choose $N_{\mathrm{sub}}$:

$$g(N_{\mathrm{sub}}) = \frac{\mathrm{Dist}(N_{\mathrm{sub}})}{N_{\mathrm{sub}} * \log(N_{\mathrm{sub}} + 1)},$$

where $\mathrm{Dist}(N_{\mathrm{sub}}) \triangleq D_{\mathrm{KL}}(\hat{p}_{\mathrm{T}}(\tilde{Y}_{h_{\mathrm{T},N_{\mathrm{sub}}}}, \tilde{Z}_{\mathrm{T}})||\hat{p}_{\mathrm{S}}(\tilde{X}_{h_{\mathrm{S},N_{\mathrm{sub}}}}, \tilde{Z}_{\mathrm{S}}))$ is obtained when we have $C_Y$, $C_X$, $C_Z$, $h_{\mathrm{T},N_{\mathrm{sub}}}$ and $h_{\mathrm{S},N_{\mathrm{sub}}}$ by solving optimization problem (S.9). There is a bias that KL divergence will tend be larger when $N_{\mathrm{sub}}$ is larger, so we penalize it with $N_{\mathrm{sub}} * \log(N_{\mathrm{sub}} + 1)$. We choose $N_{\mathrm{sub}}$ that has the lowest $g(N_{\mathrm{sub}})$ where $N_{\mathrm{sub}} \in \{1, \ldots, \min\{N_{\mathrm{S}}, N_{\mathrm{T}}\}\}$.

##### 1.4 Data generation in simulation

Similar to the simulation scheme designed by [3], we generate the data T, U and source data S in simulation study as follows (we fix $q = 1000$ and vary $w$, $\sigma_1$, $\sigma_2$ and $d$):

1. Generate $\mathbf{w}^{acc}$ and $\mathbf{w}^{exp}$.

$$w_{rj}^{acc} = \begin{cases} w, & r = 1, j = 1, \ldots, q(1-w); \\ & r = 2, j = q(1-w)+1, \ldots, 2q(1-w) \\ 1-w, & r = 2, j = 1, \ldots, q(1-w); \\ & r = 1, j = q(1-w)+1, \ldots, 2q(1-w). \end{cases}$$

$w_{1j}^{acc} = w_{2j}^{acc} \sim Beta(0.5, 2), j = 2q(1-w)+1, \ldots, 2q$. $w_{cj}^{exp} \sim Beta(w_{cj}^{acc}, 10), j = 1, \ldots, 2q(1-w); w_{1j}^{exp} = w_{2j}^{exp} \sim Beta(w_{1j}^{acc}, 10), j = 2q(1-w)+1, \ldots, 2q$.

2. Generate $z^{acc}$ and $z^{exp}$. The cluster labels are generated with equal probability 0.5.

3. Generate $u^{acc}$ and $\tilde{u}^{acc}$. $\tilde{u}_{ij}^{acc} \sim Bernoulli(0.5)$ if $u_{ij}^{acc} = 1$, where $u_{ij}^{acc} \sim Bernoulli(w_{cj}^{acc})$ if $z_{ic}^{acc} = 1, i = 1, \ldots, n_{\mathrm{S}}; \tilde{u}_{ij}^{acc} = 0$ otherwise.

4. Generate $u^{exp}$ and $\tilde{v}^{exp}$. $\tilde{v}_{lj}^{exp} \sim Bernoulli(0.8)$ if $u_{lj}^{exp} = 1$, where $u_{lj}^{exp} \sim Bernoulli(w_{lj}^{exp})$ if $z_{lc}^{exp} = 1; \tilde{v}_{lj}^{exp} \sim Bernoulli(0.1)$ otherwise; $l = 1, \ldots, n_{\mathrm{T}}$.

5. Generate $C$ and $G$. $C_{ij} \sim N(0, \sigma_1^2)$ if $\tilde{u}_{ij}^{acc} = 0$ and $C_{ij} \sim N(2, \sigma_1^2)$ if $\tilde{u}_{ij}^{acc} = 1$; $G_{lj} \sim N(0, \sigma_2^2)$ if $\tilde{v}_{lj}^{exp} = 0$ and $G_{lj} \sim N(2, \sigma_2^2)$ if $\tilde{v}_{lj}^{exp} = 1$.

6. Generate $\mathrm{T}^0$ and $\mathrm{S}^0$. $\mathrm{T}^0_{lj} = 1$ if $\mathrm{G}_{lj} > 0$ and $\mathrm{T}^0_{lj} = 0$ otherwise; $\mathrm{S}^0_{ij} = 1$ if $C_{ij} > 0$ and $\mathrm{S}^0_{ij} = 0$ otherwise.

7. Generate source data S and target data with linked features T. We choose the first $q$ columns of $\mathrm{S}^0$ as the source data S, i.e. $\mathrm{S}_{lj} = \mathrm{S}^0_{lj}, j = 1, \cdots, q$, and choose the first $q$ columns of $\mathrm{T}^0$ as the target data T, i.e. $\mathrm{T}_{lj} = \mathrm{T}^0_{lj}, j = 1, \cdots, q$.

8. Generate target data with unlinked features U by setting $\mathrm{U}_{lj} = \mathrm{T}^0_{lj} + N(0.5 + I_{\{l=2\}} * d, 0.16), j = q+1, \ldots, 2q$, where $d$ measures the distance between the mean of normal distribution when true label $l = 2$ and the mean of normal distribution when true label $l = 1$.

In our simulation, the difference on the steps of data generation from that in [3] is the addition of Steps 6-8, which generates binary data. $\mathrm{T}^0_{lj} = 1$ means gene $j$ is expressed in cell $l$, and $\mathrm{T}^0_{lj} = 0$ otherwise. $\mathrm{S}^0_{ij} = 1$ means the promoter region for feature $j$ is accessible in cell $i$, and $\mathrm{S}^0_{ij} = 0$ otherwise. More details on the notations and the simulation scheme is presented in [3].

### References


1. Dhillon IS, Mallela S, Modha DS. Information-theoretic co-clustering. Proceedings of the Ninth ACM SIGKDD International Conference on Knowledge Discovery and Data Mining. 2003; p. 89–98.

2. Dai WY, Yang Q, Xue GR, Yu Y. Self-taught Clustering. Proceedings of the 25th international Conference on Machine Learning. 2008;.

3. Lin ZX, Zamanighomi M, Daley T, Ma S, Wong WH. Model-Based Approach to the Joint Analysis of Single-Cell Data on Chromatin Accessibility and Gene Expression. Stat Sci. 2019;.


#### 2. Supplementary Figures

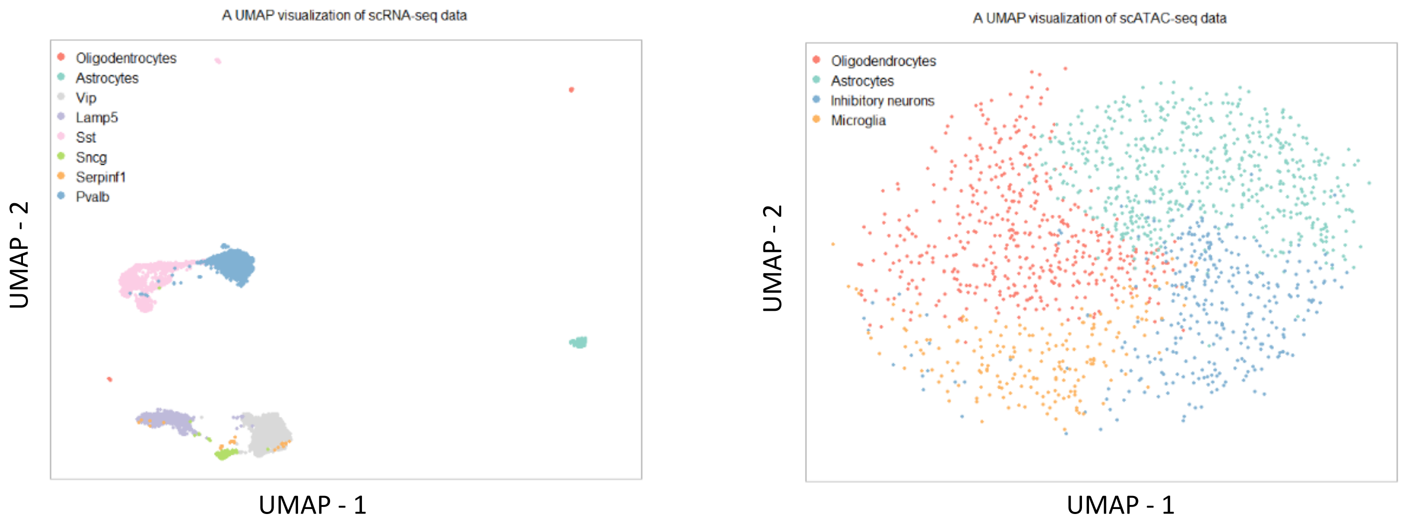


**Fig S.1.** UMAP visualization of source data (left) and target data (right) in example 1.

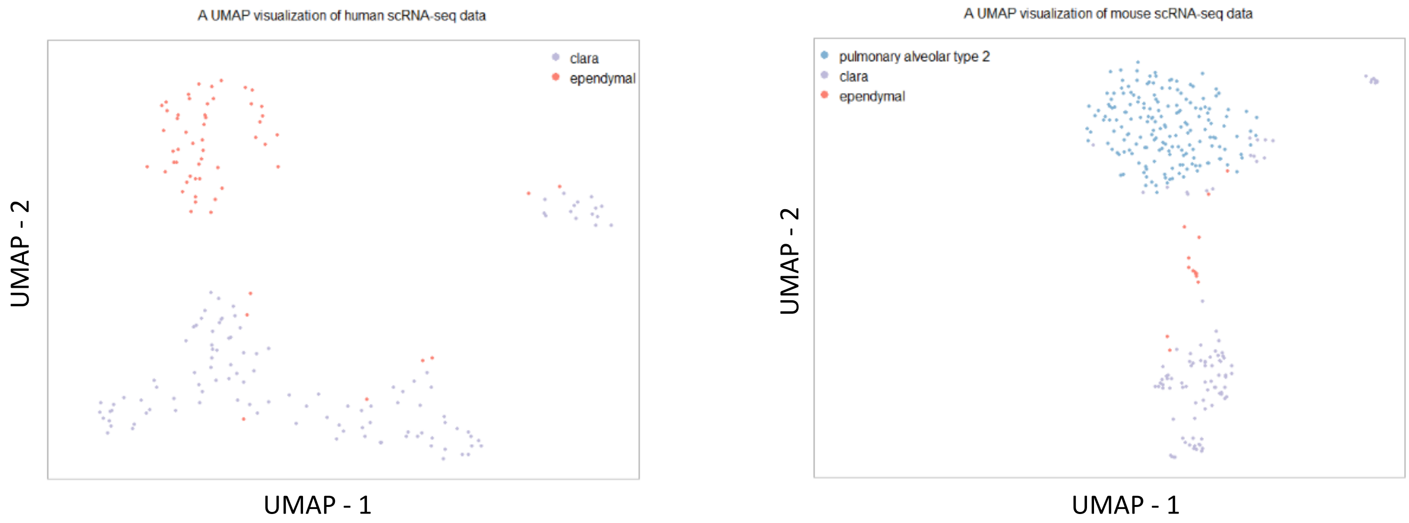


**Fig S.2.** UMAP visualization of source data (left) and target data (right) in example 2.

#### 3. Supplementary Tables

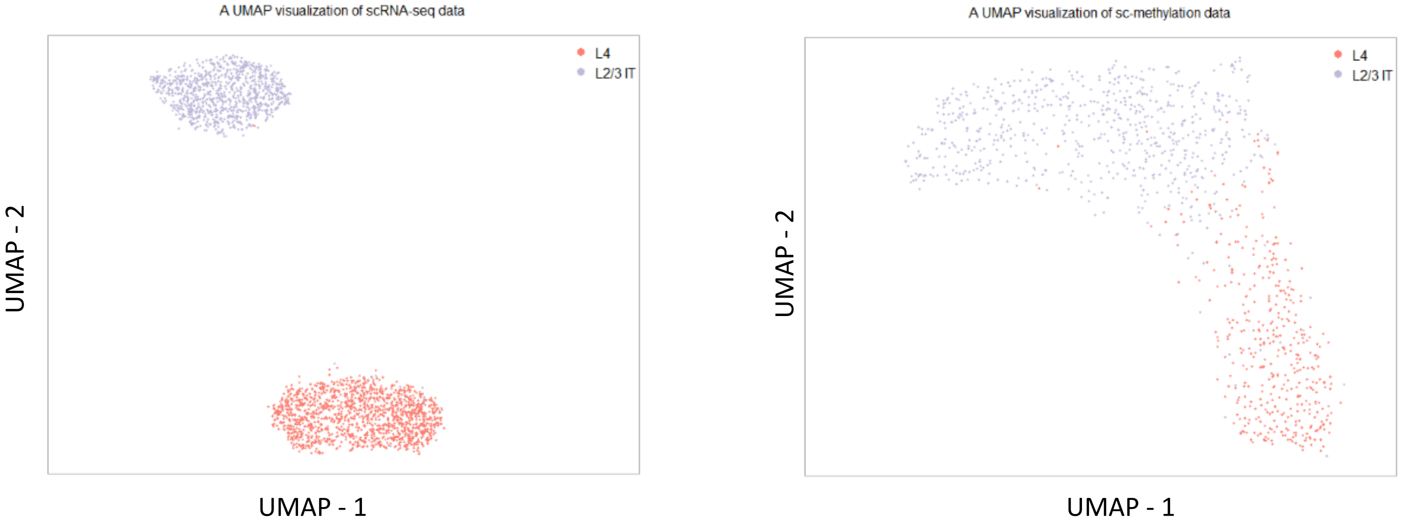


**Fig S.3.** UMAP visualization of source data (left) and target data (right) in example 3.

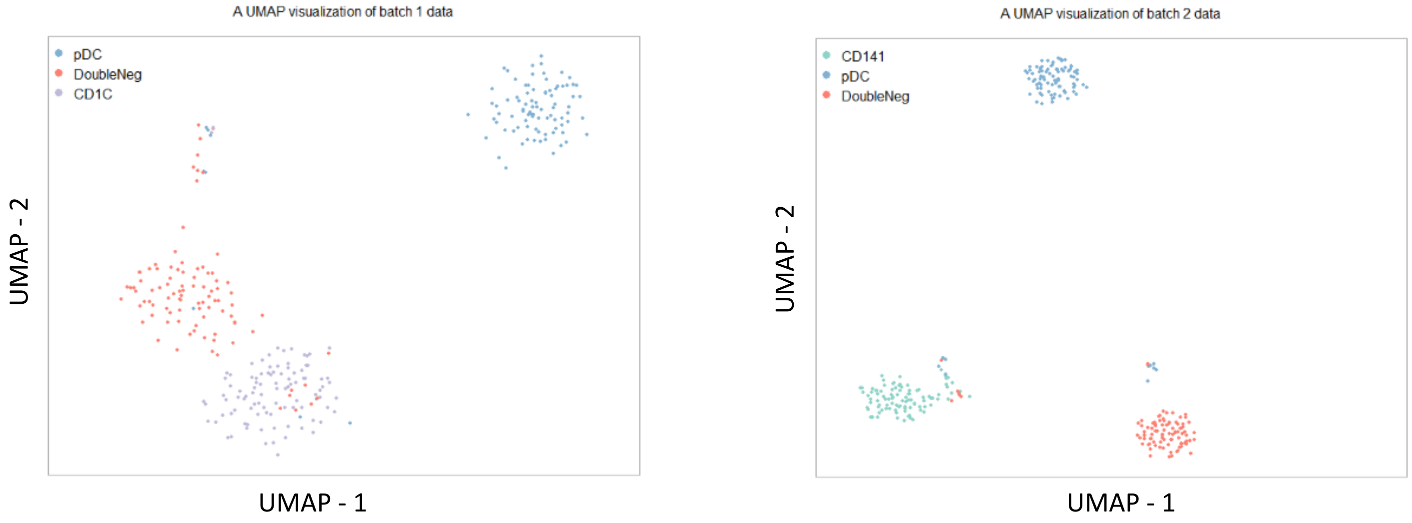


**Fig S.4.** UMAP visualization of source data (left) and target data (right) in example 4.

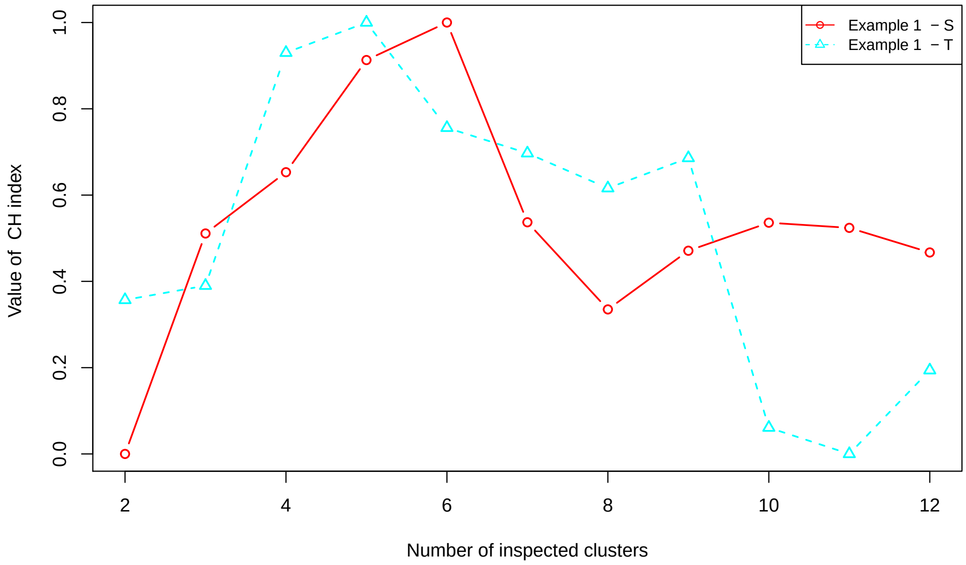


**Fig S.5.** Calinski-Harabasz evaluation on selecting the optimal number of cell clusters for the source dataset and the target dataset in example 1. The value of CH index has been standardized via minimax normalization to ensure each value being bound to between 0 and 1.

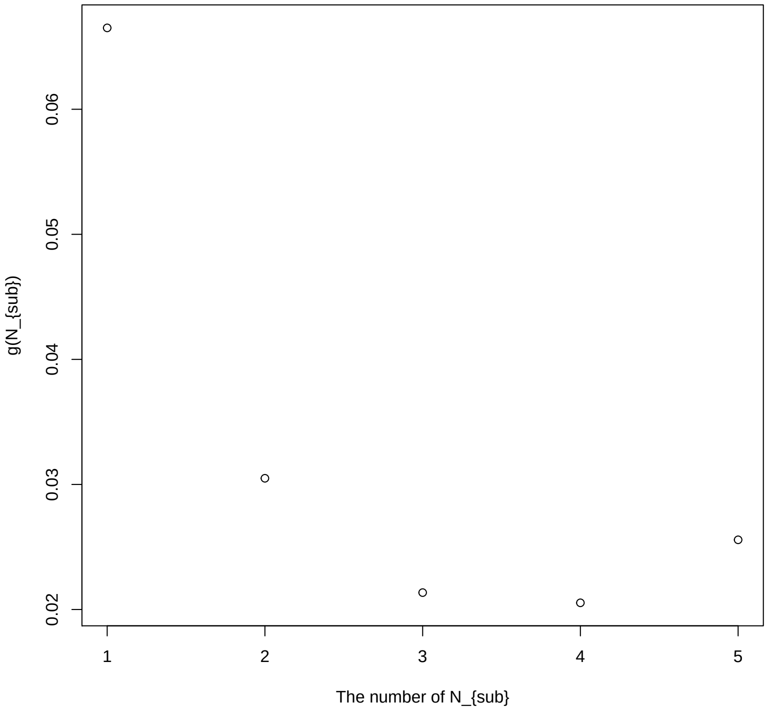


**Fig S.6.** Choose the number of $N_{\mathrm{sub}}$ in example 1.

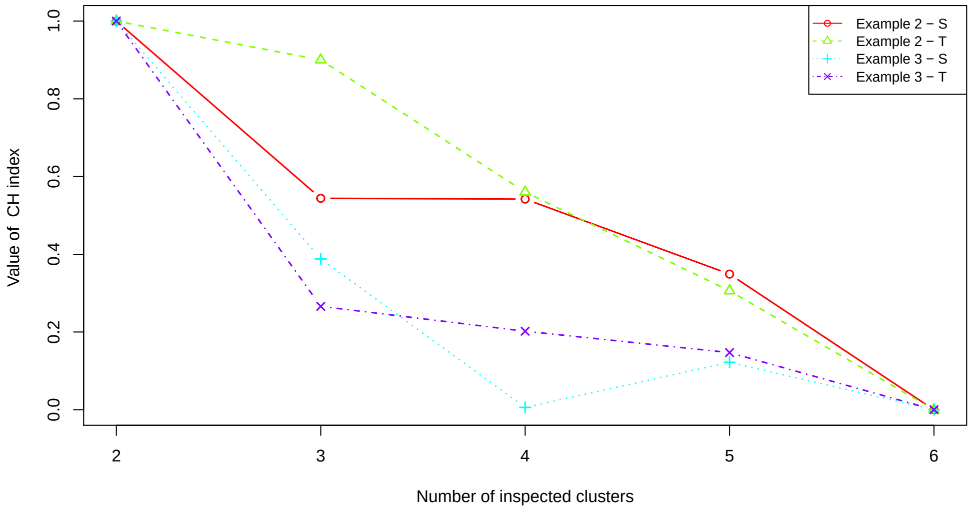


**Fig S.7.** Calinski-Harabasz evaluation on selecting the optimal number of cell clusters for the source dataset and the target dataset in examples 2 and 3. The value of CH index has been standardized via minimax normalization to ensure each value being bound to between 0 and 1.

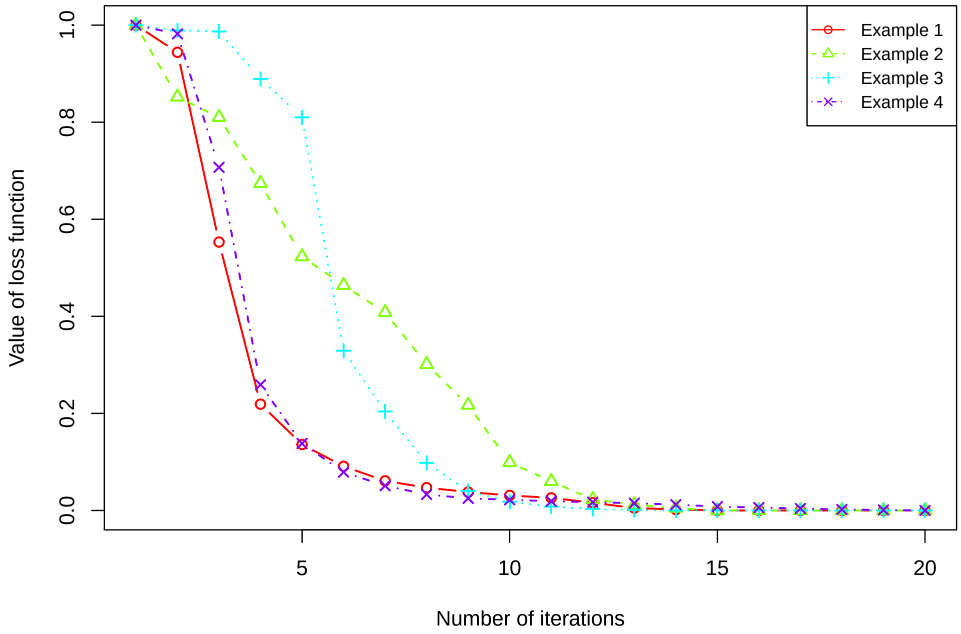


**Fig S.8.** The loss function (objective function) curves after each iteration by *couple*CoC+ in real data examples 1-4. The value of the objective function after each iteration has been standardized via minimax normalization to ensure each value being bound to between 0 and 1.

**Table S.1.** Clustering table by *couple*CoC+ in real data examples 1-4. “clu m” represents the matched cell cluster across the source data and the target data. If there is no “m” in a cell cluster label, it represents that the cluster is not matched across the two datasets, and we use “clu s” and “clu t” to represent that the cluster belongs to source data and target data, respectively.

| | | *couple*CoC+ | | | | | | | |
|---|---|---|---|---|---|---|---|---|---|
| Example 1 | | clu m1 | clu m2 | clu m3 | clu m4 | clu t5 | clu t6 | | |
| mouse scATAC-seq data | Oligodendrocytes | 1 | 11 | 446 | 0 | 0 | | | |
| (Target data, $n_{\mathrm{T}} = 1525$) | Astrocytes | 529 | 22 | 0 | 0 | 0 | | | |
| | Inhibitory neurons | 0 | 110 | 3 | 206 | 0 | | | |
| | Microglia | 0 | 0 | 0 | 0 | 197 | | | |
| | | clu m1 | clu m2 | clu m3 | clu m4 | clu s5 | clu s6 | clu s7 | clu s8 |
| mouse scRNA-seq data | Oligodentrocytes | 26 | 3 | 62 | 0 | 0 | 0 | 0 | 0 |
| (Source data, $n_{\mathrm{S}} = 6539$) | Astrocytes | 368 | 0 | 0 | 0 | 0 | 0 | 0 | 0 |
| | Vip | 0 | 13 | 0 | 0 | 0 | | 1707 | 8 |
| | Lamp5 | 0 | 49 | 0 | 0 | 0 | 0 | 28 | 1045 |
| | Sst | 0 | 153 | 0 | 1041 | 433 | 114 | 0 | 0 |
| | Sncg | 0 | 29 | 0 | 0 | 0 | 0 | 91 | 5 |
| | Serpinf1 | 0 | 0 | 0 | 0 | 0 | 0 | 19 | 8 |
| | Pvalb | 0 | 22 | 0 | 39 | 296 | 980 | 0 | 0 |

| | | *couple*CoC+ | | |
|---|---|---|---|---|
| Example 2 | | clu m1 | clu m2 | clu t3 |
| mouse scRNA-seq data | pulmonary alveolar type II | 0 | 2 | 177 |
| (Target data, $n_{\mathrm{T}} = 292$) | clara | 74 | 23 | 2 |
| | ependymal | 0 | 14 | 0 |
| | | clu m1 | clu m2 | clu s3 |
| human scRNA-seq data | clara | 110 | 3 | |
| (Source data, $n_{\mathrm{S}} = 171$) | ependymal | 0 | 58 | |

| | | *couple*CoC+ | |
|---|---|---|---|
| Example 3 | | clu m1 | clu m2 |
| mouse sc-methylation data | L4 | 26 | 386 |
| (Target data, $n_{\mathrm{T}} = 1102$) | L2/3 | 679 | 11 |
| | | clu m1 | clu m2 |
| mouse scRNA-seq data | L4 | 0 | 1401 |
| (Source data, $n_{\mathrm{S}} = 2383$) | L2/3 IT | 974 | 8 |

| | | *couple*CoC+ | | |
|---|---|---|---|---|
| Example 4 | | clu m1 | clu m2 | clu t3 |
| batch 2 scRNA-seq data | CD141 | 0 | 0 | 96 |
| (Target data, $n_{\mathrm{T}} = 288$) | pDC | 1 | 88 | 7 |
| | double negative cells | 88 | 1 | 7 |
| | | clu m1 | clu m2 | clu s3 |
| batch 1 scRNA-seq data | pDC | 1 | 91 | 4 |
| (Source data, $n_{\mathrm{T}} = 288$) | double negative cells | 89 | 0 | 7 |
| | CD1C | 2 | 0 | 94 |

**Table S.2.** Enriched functional annotation terms for gene list in the “clu 4” of linked genes in example 1 using DAVID tools. The top 10 terms are shown here. The gene list include 59 genes: PLP1, APOD, MOG, PTGDS, TGFA, NDRG1, LIMS2, CNP, MBP, MOBP, MAL, UGT8A, HAPLN2, OPALIN, RNF43, ERMN, PLAT, TNFAIP6, GM15527, PLXNB3, MAG, GJC3, EFEMP1, THSD4, GSN, FA2H, ASPA, NIPAL4, IL12A, LPAR1, GNG11, SEC14L5, BICC1, ENPEP, SPEF2, COBLL1, BCAS1, FOXS1, TSPAN2, PCOLCE, LDLRAP1, SERPINB1A, ASGR1, DDC, GAL3ST1, ST18, HEG1, CNTN2, SHROOM1, GGT6, CCDC121, GJB1, FHDC1, GM7854, INSC, SEPT4, SH3TC2, MCAM, GRB14.

| Category | Term | Count | % | Bonferroni P-value |
|---|---|---|---|---|
| GOTERM_CC_DIRECT | myelin sheath | 13 | 22.03 | 1.44E-11 |
| GOTERM_BP_DIRECT | myelination | 8 | 13.56 | 1.12E-07 |
| GOTERM_MF_DIRECT | structural constituent of myelin sheath | 4 | 6.78 | 2.51E-05 |
| UP_KEYWORDS | Glycoprotein | 24 | 40.68 | 0.00179 |
| UP_KEYWORDS | Disulfide bond | 21 | 35.59 | 0.00358 |
| GOTERM_BP_DIRECT | peripheral nervous system myelin maintenance | 3 | 5.08 | 0.0732 |
| UP_SEQ_FEATURE | glycosylation site:N-linked (GlcNAc...) | 23 | 38.98 | 0.0501 |
| UP_SEQ_FEATURE | topological domain:Extracellular | 16 | 27.12 | 0.302 |
| GOTERM_CC_DIRECT | extracellular exosome | 17 | 28.81 | 0.172 |
| UP_SEQ_FEATURE | lipid moiety-binding region:S-palmitoyl cysteine | 5 | 8.47 | 0.337 |

**Table S.3.** Enriched functional annotation terms for gene list in the “clu 6” of linked genes in example 1 using DAVID tools. The top 10 terms are shown here. The gene list include 198 genes: HCK, CXCR3, GCNT1, SLA, A630033H20RIK, TMEM173, UPK1B, P2RX7, BC035044, ITPR3, IRGM2, HMHA1, PTPN6, PIK3R5, FCGR1, RASAL3, HPGDS, HAVCR2, H2-Q7, CASP8, SAMSN1, MS4A6B, CD14, ITGAM, HCLS1, LST1, RENBP, TRAF3IP3, GPR183, SLC7A7, ALOX5, EPHA2, WHRN, IRF1, CD33, MYO1F, ENG, I830077J02RIK, CLEC4A3, RHOBTB1, FLT3, BCL2A1B, DUSP27, P2RY6, TNFRSF14, BATF, SLC40A1, FGD2, MFNG, TEC, DOCK8, LYN, HPGD, MRC1, CCL2, HLX, RPS6KA1, KLHL6, 1810011H11RIK, HVCN1, CCL9, CTSC, ANGPTL7, LAIR1, CCDC63, SUSD3, SLFN8, RHBDF2, TGFB1, IL13RA1, CEACAM1, RNASE4, AIF1, POU2F2, EDN1, GBP7, ART3, ADRB2, C430049B03RIK, CYSLTR1, CCL4, H2-T23, IGTP, PLIN2, CD300A, LRRK1, CD68, SLC11A1, CLEC4A2, C3AR1, GNA15, LRP5, CD37, VWF, H2-OA, NFAM1, CD52, RGS1, CSF3R, FLI1, PTPRC, CD86, CORT, LRMP, ST3GAL6, TLR7, LGALS9, IRS3, STAB1, GBP2, GIMAP5, TRIM47, GM12250, CD84, UGT1A7C, CCL3, TNFAIP8L2, NCF4, H2-Q6, LTC4S, CD48, SRGN, LY86, HHEX, PLD4, ALDH1A2, PSMB8, P2RY13, GLIPR1, IL10RA, VAV1, ECM1, BIN2, UCP2, PRKCH, CCL6, GGT5, PRDM1, GIMAP9, LPCAT2, GRAP, NCF1, MSN, ALOX8, LAPTM5, CX3CR1, GPR34, ABCA9, CCR5, KDR, ITGB2, FCGR2B, FILIP1L, GGTA1, KLF2, IFIT1, TMEM119, ECSCR, CSF1R, DLL4, P2RY12, FCRLS, CTSH, CD53, SLFN5, AIM2, TREM2, SLC39A8, FCGR3, XKRX, PECAM1, FAM114A1, ZC3HAV1, CFH, F11R, CTSS, BC028528, GBP9, ARHGDIB, SIGLECH, C1QA, TYROBP, SELPLG, SP100, ACVRL1, GIMAP6, FLT1, TAGLN2, SERPINF1, C1QC, ANXA3, C1QB, FCER1G, CACNA1S, IRF5, H2-EB1, PYCARD.

| Category | Term | Count | % | Bonferroni P-value |
|---|---|---|---|---|
| UP_KEYWORDS | Immunity | 35 | 17.68 | 8.27E-22 |
| GOTERM_BP_DIRECT | immune system process | 34 | 17.17 | 3.00E-19 |
| UP_KEYWORDS | Disulfide bond | 81 | 40.91 | 5.02E-19 |
| GOTERM_BP_DIRECT | inflammatory response | 29 | 14.65 | 2.81E-15 |
| UP_SEQ_FEATURE | topological domain:Extracellular | 69 | 34.85 | 2.15E-15 |
| UP_KEYWORDS | Glycoprotein | 84 | 42.42 | 1.79E-15 |
| UP_KEYWORDS | Innate immunity | 23 | 11.62 | 2.10E-14 |
| UP_SEQ_FEATURE | topological domain:Cytoplasmic | 75 | 37.88 | 3.54E-13 |
| UP_SEQ_FEATURE | disulfide bond | 69 | 34.85 | 7.79E-13 |
| GOTERM_CC_DIRECT | membrane | 121 | 61.11 | 1.72E-12 |

**Table S.4.** Summary of the computation time by classical clustering methods SC3 and SIMLR for scRNA-seq data in examples 1-3 and by *couple*CoC+ for the combination of source data and target data in examples 1-4. The algorithm *couple*CoC+ runs until convergence (15 iterations) by MATLAB R2019b - academic use. SC3 and SIMLR run in default iterations in Rstudio (Version 1.2.5033) by the downloaded R packages. All of these algorithms are run in Windows 10 Enterprise (Version 1909) with the Processor: Intel(R) Core(TM)i7-9700 CPU 3.00GHz and with 16.0 GB installed RAM.

| Examples | Clustering methods | | |
|---|---|---|---|
| | SC3 | SIMLR | *couple*CoC+ (S+T+U) |
| Example 1 ($n_{\mathrm{S}} + n_{\mathrm{T}} = 8064$) | 20.52(mins) | 55.50(mins) | 28.20(mins) |
| Example 2 ($n_{\mathrm{S}} + n_{\mathrm{T}} = 463$) | 62.99(s) | 32.35(s) | 27.82(s) |
| Example 3 ($n_{\mathrm{S}} + n_{\mathrm{T}} = 3485$) | 2.31(mins) | 35.15(mins) | 7.98(mins) |
| Example 4 ($n_{\mathrm{S}} + n_{\mathrm{T}} = 576$) | - | - | 49.72(s) |
